# Data-driven insights into the association between oxidative stress and calcium-regulating proteins in cardiovascular disease

**DOI:** 10.1101/2024.10.03.616559

**Authors:** Namuna Panday, Dibakar Sigdel, Irsyad Adam, Joseph Ramirez, Aarushi Verma, Anirudh Eranki, Wei Wang, Ding Wang, Peipei Ping

## Abstract

A growing body of biomedical literature suggests a bidirectional regulatory relationship between cardiac calcium (Ca^2+^)-regulating proteins and reactive oxygen species (ROS), which is integral to the pathogenesis of various cardiac disorders via oxidative stress signaling. To address the challenge of finding hidden connections within the growing volume of biomedical research, we developed a data science pipeline for efficient data extraction, transformation, and loading. Employing the CaseOLAP (Context-Aware Semantic Analytic Processing) algorithm, our pipeline quantifies interactions between 128 human cardiomyocyte Ca^2+^-regulating proteins and eight cardiovascular disease (CVD) categories. Our machine learning analysis of CaseOLAP scores reveals that the molecular interfaces of Ca^2+^-regulating proteins uniquely associate with cardiac arrhythmias and diseases of the cardiac conduction system, distinguishing them from other CVDs. Additionally, a knowledge graph analysis identified 59 of the 128 Ca^2+^-regulating proteins as involved in OS-related cardiac diseases, with cardiomyopathy emerging as the predominant category. By leveraging a link prediction algorithm, our research illuminates interactions between Ca^2+^-regulating proteins, OS, and CVDs. The insights gained from our study provide a deeper understanding of the molecular interplay between cardiac ROS and Ca^2+^-regulating proteins in the context of CVDs. Such understanding is essential for the innovation and development of targeted therapeutic strategies.

## 1. Introduction

Calcium (Ca^2+^) plays a pivotal role in various biological systems, serving as an essential messenger in numerous cellular processes [1–5]. One such critical system is the cardiovascular system, where a network of Ca^2+^-regulating proteins is involved for cardiac functionality. Proteins such as L-type Calcium Channel (LTCC), Ryanodine Receptor 2 (RyR2), and troponin C—a myofilament protein—engage in a tightly coordinated series of steps that govern Ca^2+^ dynamics [6–11]. These dynamics are crucial for the excitation-contraction cycle (ECC), the process that allows the heart to contract and relax efficiently. In addition to their role in the ECC, Ca^2+^-regulating proteins also have vital roles in cellular energetics. Mitochondrial Ca^2+-^regulating proteins —including the mitochondrial Ca^2+^ uniporter (MCU), mitochondrial sodium/calcium exchanger protein (NCLX), and mitochondrial calcium uptake 1 and 2 (MCU1 and MCU2)—help maintain Ca^2+^ homeostasis in the mitochondria. This is essential not only for normal Adenosine Triphosphate (ATP) generation but also for the regulation of Reactive Oxygen Species (ROS), highlighting the interconnectedness of Ca^2+^-regulatory protein networks in various biomolecular processes [12–14].

The intricate relationship between Ca^2+^ and ROS is worth noting. These two agents mutually regulate one another in cellular dynamics. Elevated Ca^2+^ concentrations within a cell can lead to an overproduction of ROS, inducing oxidative stress. This stress is essentially a disruptive imbalance between oxidants, such as ROS, and reductants like antioxidants. This abnormal surge in ROS levels is notorious for causing damage to proteins, lipids, and DNA—a chain of events that ultimately causes cell death [15–21]. Conversely, an abundance of ROS can have profound impacts on cellular Ca^2+^ dynamics. This is evident as ROS can remodel Ca^2+^-regulating ion channels, disrupt associated pathways, and alter the functions of regulatory proteins [22–24]. Such perturbations in Ca^2+^ dynamics are linked to CVDs like arrhythmia, heart failure, and contractile dysfunction [22, 25–27]. Given the significant overlap in the literature about Ca^2+^-ROS dynamics and CVDs, it becomes vital to systematically explore these interconnections.

To address this need, we utilized advanced text-mining pipelines, specifically employing the CaseOLAP pipeline [28, 29]. This tool allowed us to categorize publications into eight distinct CVD categories and score the relevance of 128 cardiac Ca^2+^-regulating proteins within these categories. The derived scores were based on two critical components: ‘popularity’, measuring the frequency of a given protein in one category versus others in the same category, and ‘distinctiveness’, assessing the frequency of the target protein in one category compared to alternate categories. Impressively, these scores remain robust even among class imbalances, like varied document counts in CVD categories. A higher score signifies a stronger protein-disease association.

Furthering our analytical endeavors, we formulated a knowledge graph (KG) [30, 31]. This KG combined the scored protein-disease associations with other associated datasets, such as proteins, CVD and OS MeSH descriptors, reference articles and molecular pathways. The KG analysis provided a comprehensive view, highlighting associations between cardiac Ca^2+^-regulating proteins, CVDs, and OS signaling networks. By revealing these intricate relationships, we aspire that our comprehensive approach will pave the way for future research, potentially guiding the discovery of novel drug targets and therapeutic avenues for CVDs.

## 2. Materials and Methods

This study is designed to elucidate the shared molecular mechanisms between Ca^2+^-regulating proteins and oxidative stress molecules in relation to eight CVD categories as outlined in Table 1A. To achieve this objective, we have established a comprehensive platform that integrates text mining with knowledge graph (KG) analysis.

**Table 1A.** Document statistics per CVD categories. Documents collection statistics in each CVD category and the OS-CVD interface: We collected a total of 1,197,530 unique documents, all studying at least one CVD. Each CVD category is broken down by name, abbreviation, associated MeSH tree numbers, the number of documents studying the CVD category, and the number of documents also studying OS.

| CVD Category | Abbreviation | Major Root Nodes (MeSH) | No. of CVD publications collected | No of OS related publications within CVD |
| --- | --- | --- | --- | --- |
| Cardiomyopathies and Heart Failure | CM | C14.280.238, C14.280.434 | 247,436 | 34,063 |
| Arrhythmias, Cardiac | ARR | C14.280.067 | 239,060 | 15,960 |
| Heart Defects, Congenital | CHD | C14.280.400 | 154,992 | 7,183 |
| Heart Valve Diseases | VD | C14.280.484 | 137,197 | 4,127 |
| Myocardial Ischemia | IHD | C14.280.647 | 473,233 | 50,435 |
| Cardiac Conduction System | CCS | C14.280.123 | 100,841 | 6,613 |
| Ventricular Outflow Obstruction | VOO | C14.280.955 | 42,942 | 1,003 |
| Other Heart Diseases (Cardiomegaly, Endocarditis, Heart Arrest, Heart Rupture, Ventricular Dysfunction, Heart Neoplasms, Pericarditis) | OHD | C14.280.195, C14.280.282, C14.280.383, C14.280.470, C14.280.945, C14.280.459, C14.280.720 | 215,416 | 17,295 |

### 2.1. Data collection

#### 2.1.1. Document collection

Initially, we curated a comprehensive corpus of biomedical literature until October 2023, focusing on eight CVD categories and their association with oxidative stress. A total of 1,197,530 unique documents were identified that explored CVDs, with 102,807 of these also addressing oxidative stress (refer to **Table 1A**). The selection of documents was guided by relevant CVD and OS Medical Subject Headings (MeSH) descriptors, which are systematically arranged in a hierarchical framework [32]. This structure facilitated the retrieval of publications across various levels of specificity concerning the CVD categories and oxidative stress.

#### 2.1.2. CVD-MeSH collection

In this study, we employed eight predefined categories of CVDs as outlined in **Table 1A** [33, 34]. We compiled associated MeSH for these CVD categories, including both root nodes and their descendant entries, from the MeSH Library of the National Library of Medicine (NLM) database. This comprehensive retrieval resulted in a total of 176 unique CVD MeSH descriptors. These descriptors are detailed in the supplementary material, presented in both a tree structure visualization and a table (**Table S1**) format.

#### 2.1.3. Oxidative Stress (OS)-MeSH collection

Oxidative stress (OS) is an imbalance between oxidants, such as ROS, and reductants, such as antioxidants, leading to an abnormal increase in ROS levels. In our study, we manually collected OS-relevant molecules from the literature and mapped them to the corresponding MeSH descriptors using the NLM MeSH Library. A total of 75 OS-MeSH descriptors were utilized in this study. We categorized these OS MeSH descriptors into three distinct phases: Initiation of OS (IOS), Modulation of OS (MOS), and Outcome of OS (OOS). Each category is briefly described in the following sections. A detailed classification of OS MeSH descriptors is available in the supplemental material, presented in both a tree structure visualization and a table (**Table S2**) format.

##### 2.1.3.1. Initiation of OS (IOS)

IOS includes all OS events involved in producing free radical and non-radical species. This phase incorporates three subcategories of chemically reactive species: ROS which contains oxygen, reactive nitrogen species (RNS) which are derived from nitric oxide, and reactive aldehydes (RA) which are organic compounds with a carbonyl functional group. IOS includes 12 molecules, enzymes, or proteins.

##### 2.1.3.2. Modulation of OS (MOS)

MOS includes events involving the OS process and progression. This phase incorporates four subcategories: redox metabolites which result from oxidation/reduction reactions removing oxidative radicals, antioxidants which contain compounds that inhibit or eliminate oxidation and free radical release, antioxidant enzymes which contain enzymes that catalyze free radical decomposition, and redox regulating proteins which contain proteins involved in redox signaling. MOS includes 59 molecules, enzymes, or proteins.

##### 2.1.3.3. Outcomes of OS (OOS)

OOS includes OS events involved in the downstream consequences and products of OS. This phase incorporates four subcategories: protein oxidation which contains protein products following reactions with ROS, lipid peroxidation products which contain biochemical products of lipid oxidation, oxidative DNA damage which contains oxidative lesions in DNA, and nitrative DNA damage which contains nitrative lesions in DNA. OOS includes 6 molecules, enzymes, or proteins.

#### 2.1.4. Assembly of cardiac ca^2+^-regulating protein list

Initially, we undertook an advanced search on UniProt using targeted keywords like “ Ca^2+^ ion channels,” “heart,” and “human.” This search yielded 105 reviewed proteins linked to cardiac Ca^2+^ dynamics. Further, manual curation was conducted by our team to focus exclusively on proteins that hold functional significance and play a direct, pivotal role in ensuring Ca^2+^ homeostasis in the cardiac cell based on the GO terms, cardiac proteome data and relevant literature search [35]. After rigorous curation, such as excluding certain tissue-specific proteins, we compiled 128 Ca^2+^-regulating proteins, each associated with a distinct UniProt ID. The comprehensive classification of these proteins is presented in the Supplemental material, both in a tree structure visualization and a tabulated format (**Table S3**), detailing protein names alongside their UniProt IDs.

#### 2.1.5. Pathway collection

All pathways associated with the 128 Ca^2+^-regulating proteins were extracted from Reactome knowledgebase.

### 2.2. Workflow design

We implemented a text mining and knowledge graph platform to better understand the underlying molecular mechanisms of Ca^2+^-regulating proteins involved in 8 CVDs and OS. We executed the CaseOLAP score to reveal the association between Ca^2+^-regulating proteins and 8 CVD categories. We analyzed those protein-disease association scores with unsupervised machine learning techniques (PCA and hierarchical clustering) [36, 37] to gain further insight into the shared roles of proteins. We then constructed a KG graph by incorporating scored proteins, their pathways, CVD and OS MeSH descriptors and PubMed documents related to CVDs and OS. The workflow (**Figure 1**) illustrates this process. We further explored the knowledge graph using smart queries and a link prediction algorithm to reveal and propose hidden relationships between the Ca^2+^-regulating proteins, OS, and CVDs.

**Figure 1.**
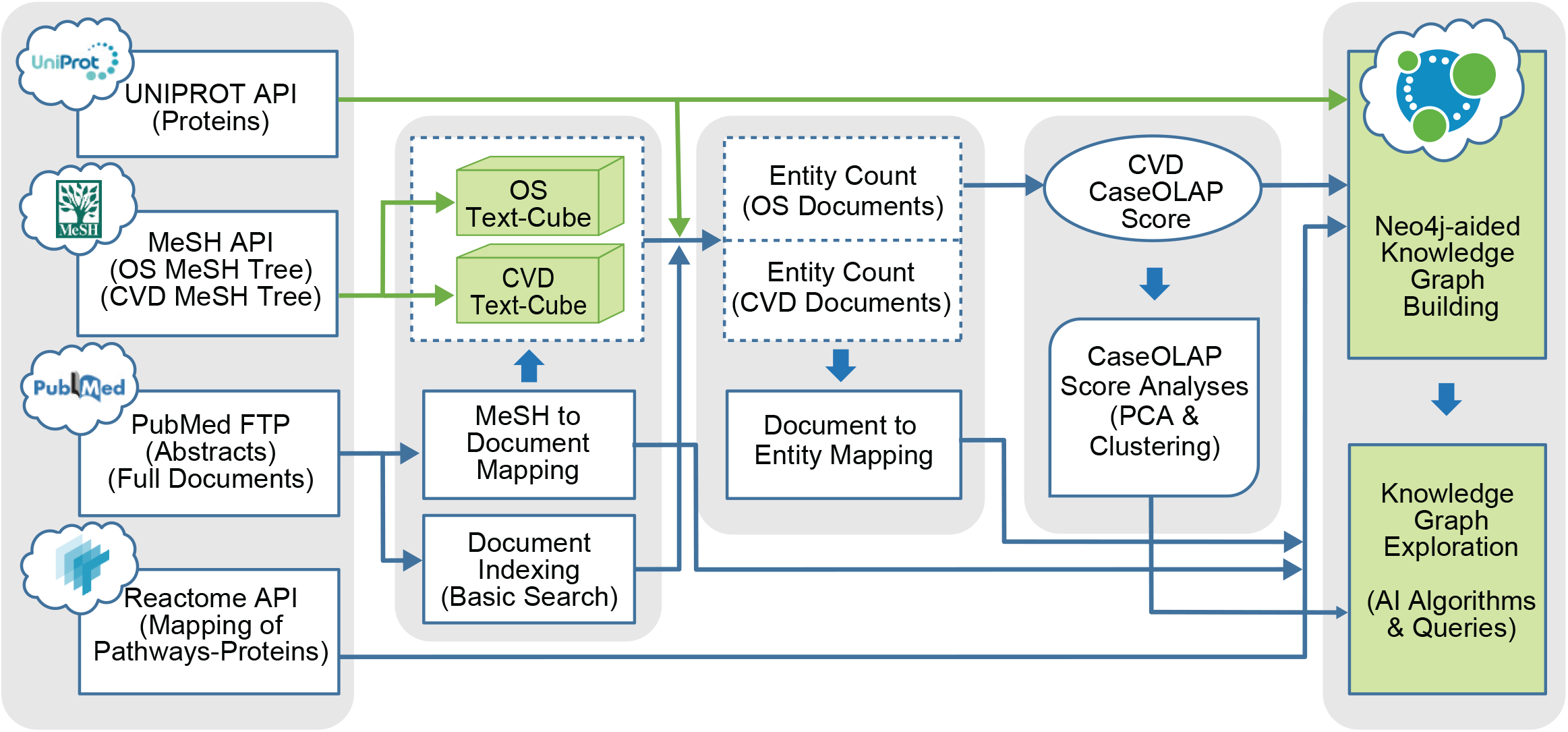
Overview of the Workflow. This workflow illustrates the process of extracting, transforming, loading, and analyzing the relevant data sources. The leftmost column represents the data sources: UniProt for the Ca^2+^-regulating proteins, MeSH for oxidative stress (OS) concepts and 8 categories of CVDs, PubMed (https://pubmed.ncbi.nlm.nih.gov/) for documents, and Reactome (https://reactome.org/) for relevant pathways. Two text cubes were assembled for collections of relevant documents studying OS and CVDs respectively. CaseOLAP scores were computed to quantify the relevance with respect to the CVD categories for every relevant protein. These information were integrated via the construction of a knowledge graph along with documents, MeSH descriptors, and pathways.

### 2.3. Knowledge Graph Construction

We constructed a heterogeneous KG from our protein-disease association scores and other biomedical data sources (UniProt, PubMed, MeSH, and Reactome). Our KG included four node types (protein, document, MeSH, and pathway) and three edge types (document-Assigns-MeSH, document-Mentions-protein, and pathway-Contains-protein). **Figure 2** shows the KG schema and **Table 1B** and **1C** list node and edge statistics, respectively.

**Table 1B.** Node statistics. The total number of nodes and edges in our KG are provided in the tables.

| Nodes | Total unique nodes |
| --- | --- |
| Document Nodes | 1,197,530 |
| MeSH Nodes | 251 (75 are OS, 176 are CVD) |
| Protein Nodes | 128 |
| Pathway Nodes | 496 |

**Table 1C.**
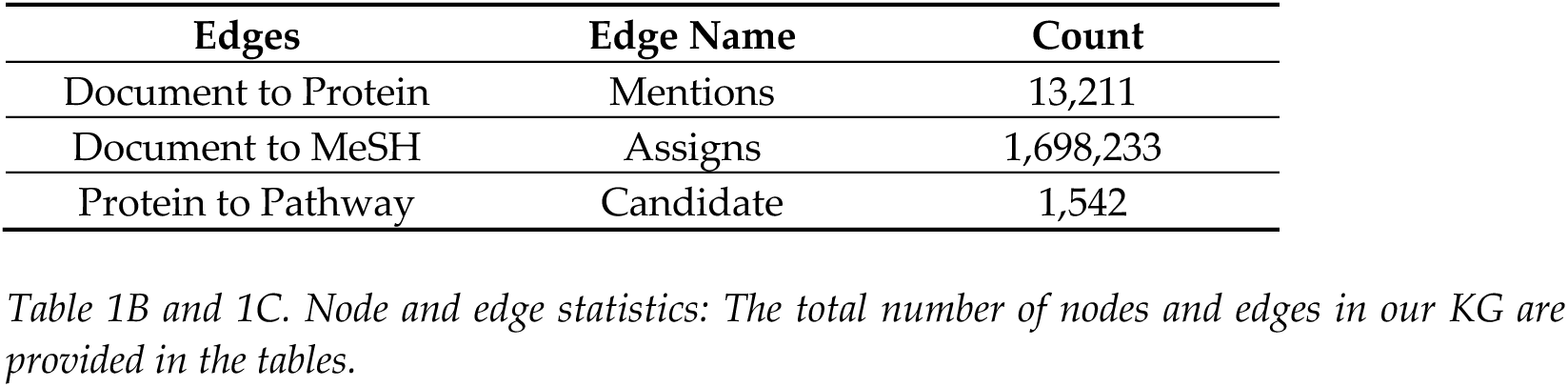
Edge statistics. The total number of nodes and edges in our KG are provided in the tables.

**Figure 2.**
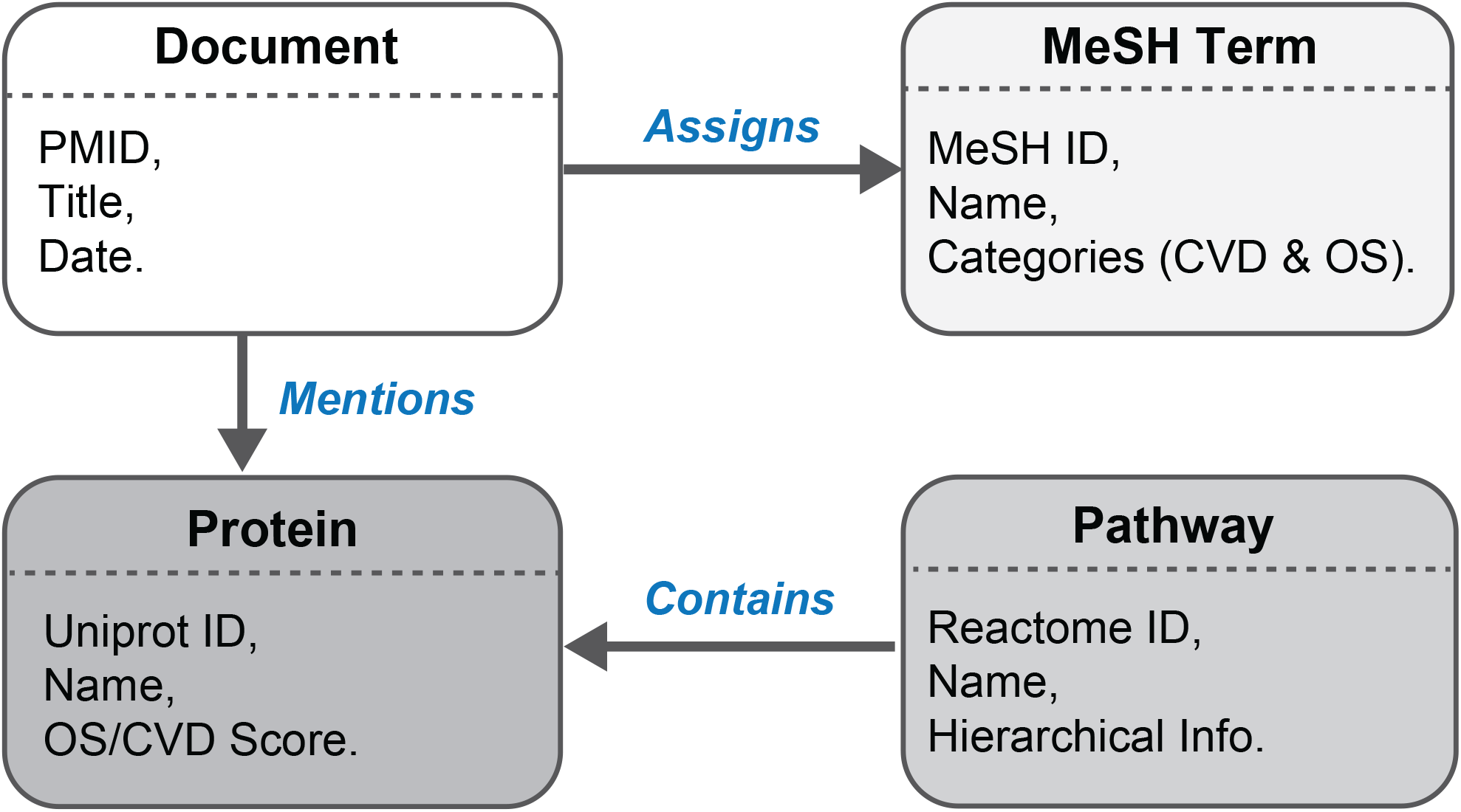
Knowledge Graph Data Schema. This data schema represents the Knowledge Graph (KG) structure, its node, and edge types. There are 4 nodes (MeSH, Document, Protein, and Pathway), as well as 3 edges (Document-Assigns-MeSH, Document-Mentions-Protein, and Pathway-Contains-Protein).

## 3. Results

### 3.1. Interaction among Ca^2+^-regulating proteins, CVD and OS categories

We systematically examined the interaction among Ca^2+^-regulating proteins, CVD, and OS categories from a quantitative perspective, as depicted in **Figure 3 (A and B)**. Specifically, **Figure 3A** elucidates the number of shared documents across distinct CVD and OS categories, while **Figure 3B** delineates the overlap of proteins across CVD and OS categories.

**Figure 3.**
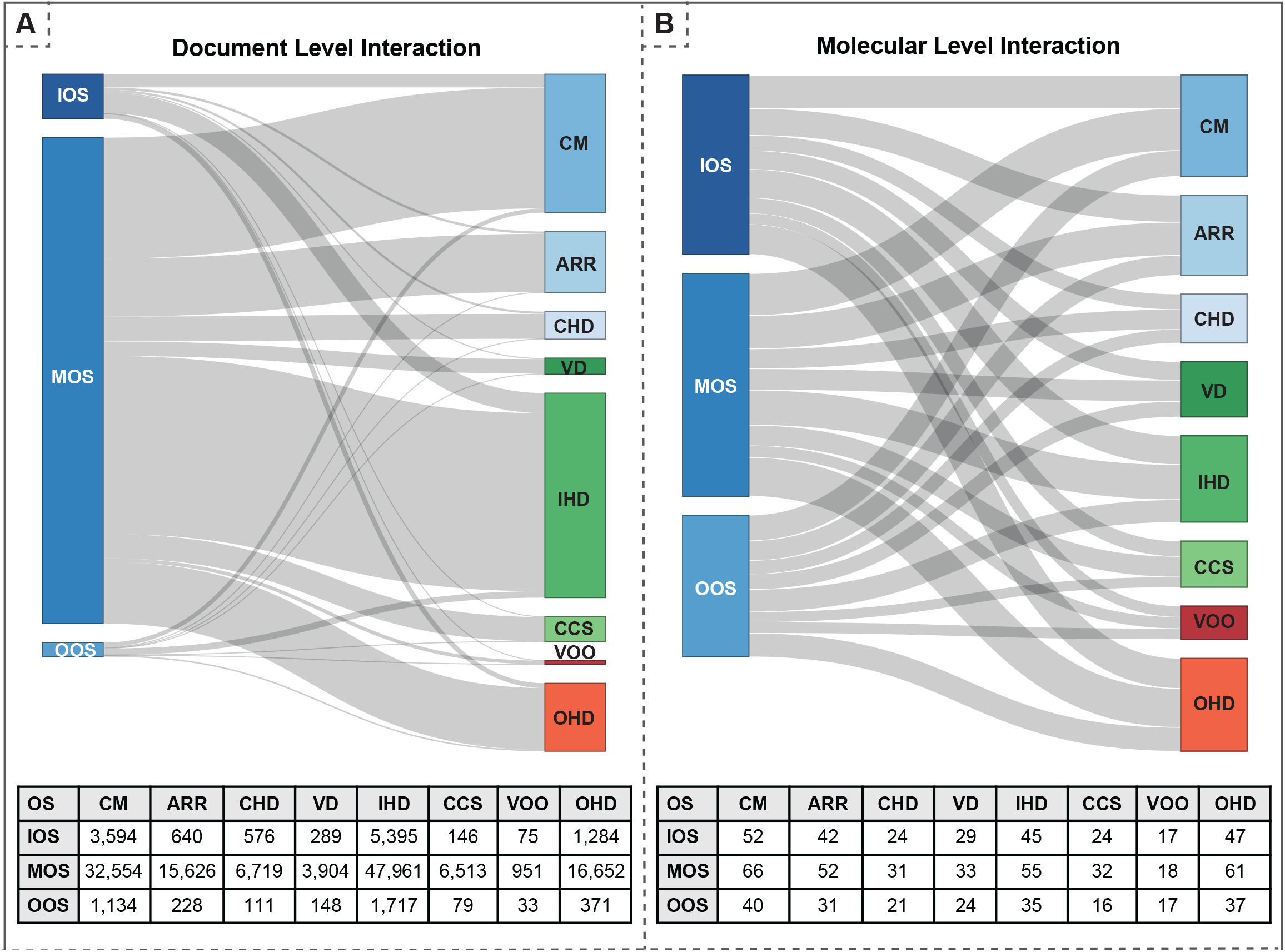
Visualization of document-level association and molecular-level interaction between OS and CVD Categories. **A.** The Top Panel is a Sankey diagram outlining the document-level OS-CVD association. The Column Height represents the total number of documents associated with each OS phases and CVD categories; the Chord Thickness is defined by the proportion of documents shared between each OS-CVD pair. An interactive version of this plot is also available online (https://caseolap.github.io/IonChannel/rcodes/doc-intersection.html). The Bottom Panel shows the detailed document counts for the Sankey diagram above. B. The Top Panel is a Sankey diagram representing the molecular-level interaction between OS and CVD catergories that are specificially relevant to Ca^2+^ regulating proteins. The Column Height represents the total number of proteins related to each OS phases and CVD categories, and the thickness of the chord represents the number of proteins shared among each OS phases and CVD categories. The interactive version is available online (https://caseolap.github.io/IonChannel/rcodes/protein-intersection.html). The bottom panel provides the numerical data of the Sankey diagram displayed above.

#### 3.1.1. Documents in OS-CVD Categories

We collected 102,807 unique documents studying CVDs and OS. A pronounced volume of publications was observed at the nexus of modulation of oxidative stress (MOS) with both ischemic heart disease (IHD) and cardiomyopathy (CM). Although the distribution of publications in pairwise OS-CVD categories was disproportionate, each OS-CVD category contained at least some publications. This encouraged us to explore the protein level interaction (e.g., proteins behind the OS-CVD associations) by identifying the proteins in each OS-CVD category.

#### 3.1.2. Proteins in OS-CVD Categories

Inspired by the interaction seen at the document level (see **Figure 3(A and C)**), we further quantified protein-level interaction among the CVD and OS categories (see **Figure 3 (B and D)**). The strength of the interaction is calculated based on the shared proteins with a non-zero CaseOLAP score. Although the documents were disproportionately represented in the OS-CVD categories, the proteins were more uniformly distributed. This suggests that, despite the disproportionate studies in the OS-CVD categories (as represented by the disproportionate number of publications), we can obtain useful biological information: Ca^2+^-regulating proteins serve biological functions in each stage of OS in each major CVD category.

### 3.2. CaseOLAP score analysis

In our analysis of the corpus, a notable 78 out of the 128 identified Ca^2+^-regulating proteins acquired a CaseOLAP score in relation to at least one CVD category. This indicates an association of these proteins with one or more of the eight delineated CVDs. A comprehensive representation of all scoring proteins across CVD categories is illustrated in **Figure 4A**. The association patterns of proteins within individual CVDs seem heterogeneous; these proteins exhibit varied relationships across the eight CVDs. Notably, cardiomyopathy (CM) is linked with 67 of the 78 scoring proteins, underscoring its significant correlation with Ca^2+^-regulating protein functionality. Similarly, ischemic heart disease (IHD) and arrhythmias (ARR) have associations with 55 and 52 proteins, respectively. By utilizing CaseOLAP scores, we further implemented dimensionality reduction and clustering methods to discern the scoring patterns of these proteins.

**Figure 4.**
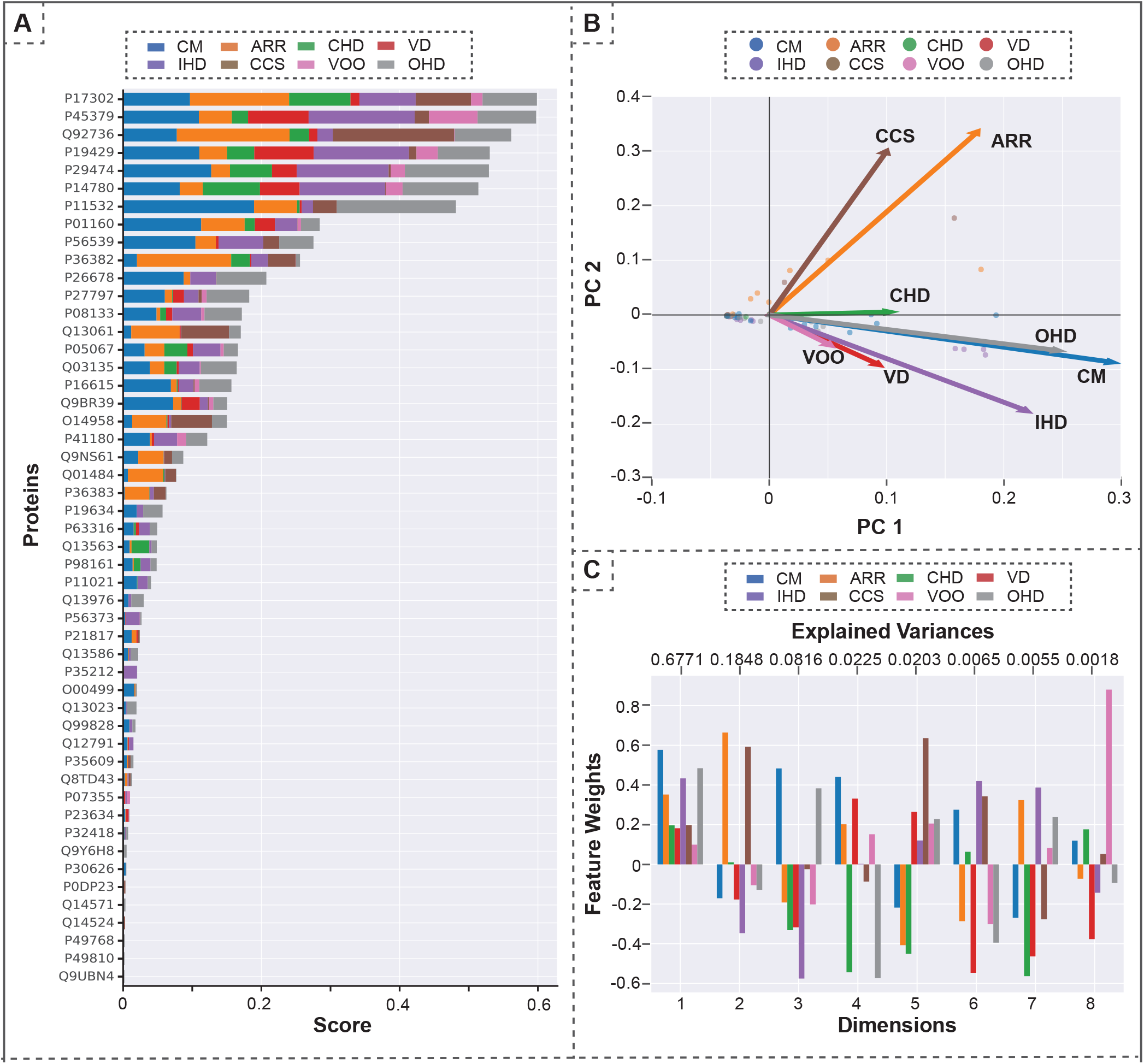
Analysis of top-scoring proteins and PCA results: Panel A demonstrates the top-scoring proteins across all CVD categories, visualized as a stacked bar chart. Panel B illustrates the results of the protein’s principal component analysis (PCA) within the 8-dimensional vector space corresponding to the CVD categories. This is subsequently projected onto a 2-dimensional plane defined by the primary principal components (PC1 and PC2). Individual dots signify distinct proteins, while the arrow-headed vectors represent the eight CVD categories in relation to PC1 and PC2. This illustration helps in comparing the behavior of the CVD categories based on the scoring proteins, providing insights into potential groupings of proteins operating via analogous molecular mechanisms. Notably, CCS and ARR emerge as distinct from the remaining CVDs. Panel C indicates the contribution of each CVD within the PCA dimensions. Considering the first two dimensions (i.e., PC1 and PC2), the first dimension is the positive superposition of all diseases, and ARR and CCS dominate the second dimension. The color bar in A, B and C represents different CVD categories.

#### 3.2.1. Principal Component Analysis (PCA)

PCA is a machine learning technique for dimensionality reduction ^35^. We utilized PCA to transform the 8-dimensional protein score vectors into a more understandable 2-dimensional space, as depicted in **Figure 4B**. Within the PCA plot, each dot symbolizes a distinct protein, whereas each arrowhead vector provides a 2D projection corresponding to a specific CVD category. A notable observation from this analysis is the distinct positioning of two CVDs, ARR and CCS, which are differentiated from the other six categories based on their respective CaseOLAP scores. Figure 4C showcases the factor loadings of the principal components. PC1 comprises a relatively even combination of all CVDs, while PC2 is dominated by CCS and ARR.

#### 3.2.2. Clustering behavior of Ca^2+^-regulating proteins

Using a Euclidean distance metric, we applied hierarchical clustering [36] to the proteins under study based on their 8-dimensional CaseOLAP scores across the 8 CVD categories. This clustering technique groups proteins with similar scores, reflecting their potential biomedical relevance within specific CVDs. Our analysis recognized distinct protein clusters corresponding to specific CVDs based on their CaseOLAP scores, as shown in **Figure 5**. For instance, proteins enclosed within the red boundary predominantly influence the ARR and CCS categories in contrast to other CVDs. Similarly, a cluster of proteins enclosed within the yellow square show a pronounced association with CM, IHD and CCS. These insights guided us for the subsequent knowledge graph analyses.

**Figure 5.**
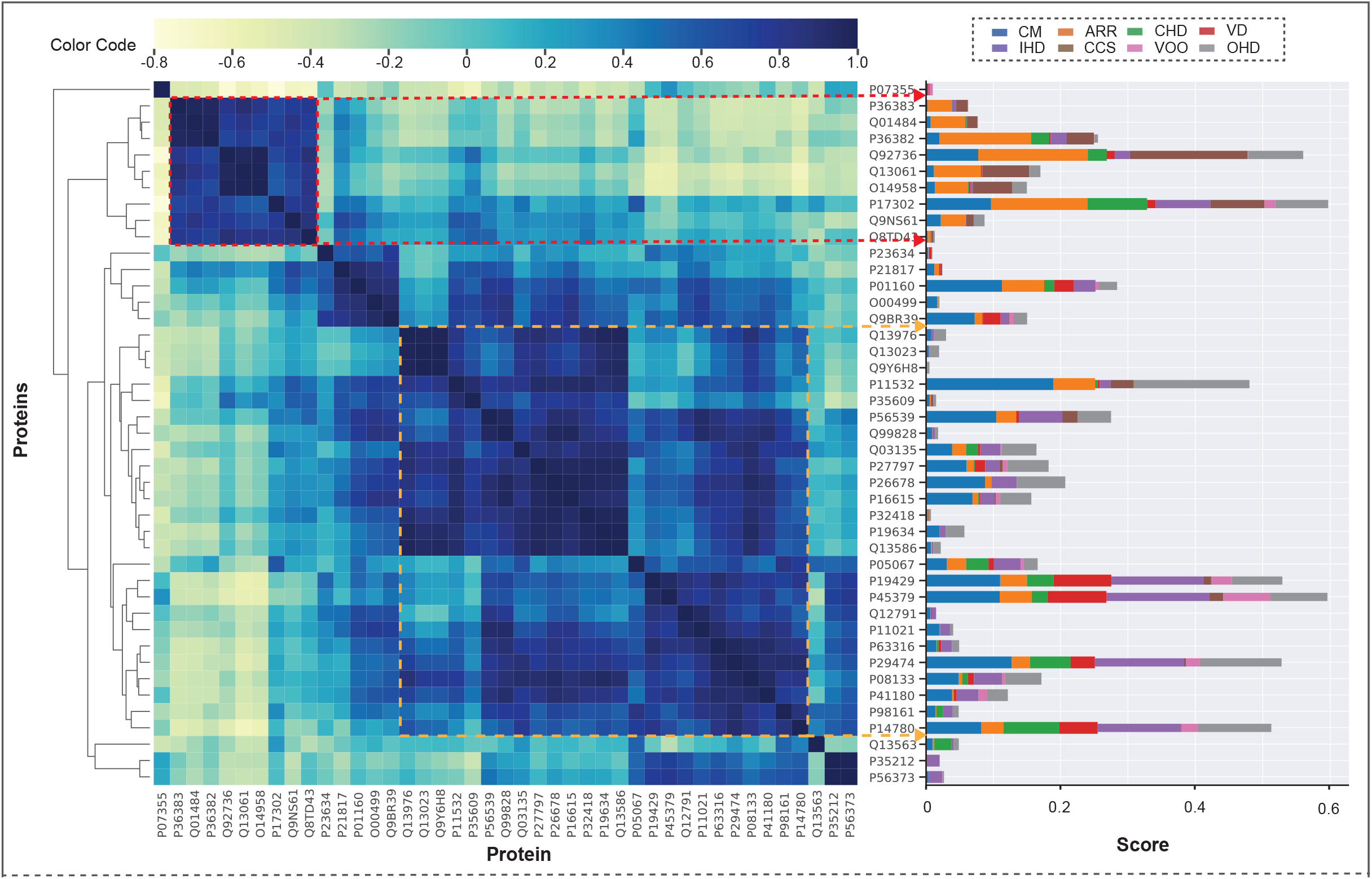
Clustering behaviors of CVD proteins. The cluster image demonstrates the hierarchical clustering of protein scores into a two-dimensional cluster plot. The cluster is formed based on the Euclidean distance metric in 8-dimensional protein vector space. The distance scores obtained are distributed in a cluster plot ranging from −0.8 to 1.0, as shown in the color legend. A darker intensity indicates closely clustered proteins. The results demonstrated two significant clusters. The cluster enclosed with a red square represents a group of proteins associated with ARR and CCS. The cluster enclosed with a yellow square represents a group of proteins related to CM, IHD and OHD. The bar plot on the right is the visualization of the CaseOLAP scores of proteins rearranged based on the clustering.

### 3.3. KG Analyses

We analyzed the KG using cypher queries and a link prediction algorithm

#### 3.3.1. KG Analysis: Queries

Implementing queries, we searched for the Ca^2+^-regulating proteins in the document corpus mentioned together with OS molecules. Our findings reveal that 59 out of the 128 examined Ca^2+^-regulating proteins demonstrate associations with at least one oxidative stress molecule and a CVD. Notably, these 59 proteins display a diverse range of CaseOLAP scores across the different categories of cardiovascular diseases. Of particular interest, 55 of these proteins are involved in cardiomyopathy, the most prevalent disease category linked to calcium regulation and oxidative stress interplay. Ischemic heart disease follows closely, with 50 proteins demonstrating associations, underscoring it as the second most prevalent condition in this context. We have provided a comprehensive breakdown of the interconnections between Ca^2+^-regulating proteins and oxidative stress molecules across each cardiovascular disease category in supplemental **Table S4**.

To further dissect the molecular mechanism of these associations, we delved into our knowledge graph, employing cypher queries to elucidate shared oxidative stress molecules and pathways that cluster within the ARR and CCS disease categories. The following subsections present the most relevant OS molecules and pathways corresponding to those proteins.

##### 3.3.1.1. Significant OS molecules in ARR and CCS cluster

In our analysis, we have identified a pivotal interplay between specific OS molecules and Ca^2+^-regulating proteins within the cluster of ARR and CCS diseases. Utilizing cypher queries within our knowledge graph, we systematically isolated proteins that co-occur with OS molecules within the same literature sources. This allowed us to calculate an average CaseOLAP score for these OS molecules across each CVD category, as defined by Equation 1:

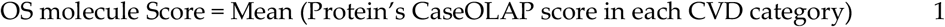

The findings are visually represented through an interactive sunburst chart (**Figure 6(A)**), which displays all the significant OS molecules in an ARR-CCS cluster based on the hierarchical relationship of OS molecules through the series of circles moving outwards according to their hierarchy. The inner, middle and outer circles represent the category, sub-category and individual molecules, respectively. The significant OS molecules are at the outermost circumference, where the thickness of the arc represents their collective scores in ARR and CCS categories. Circumference of concentric inner cords is calculated based on the total score of their descendants.

**Figure 6.**
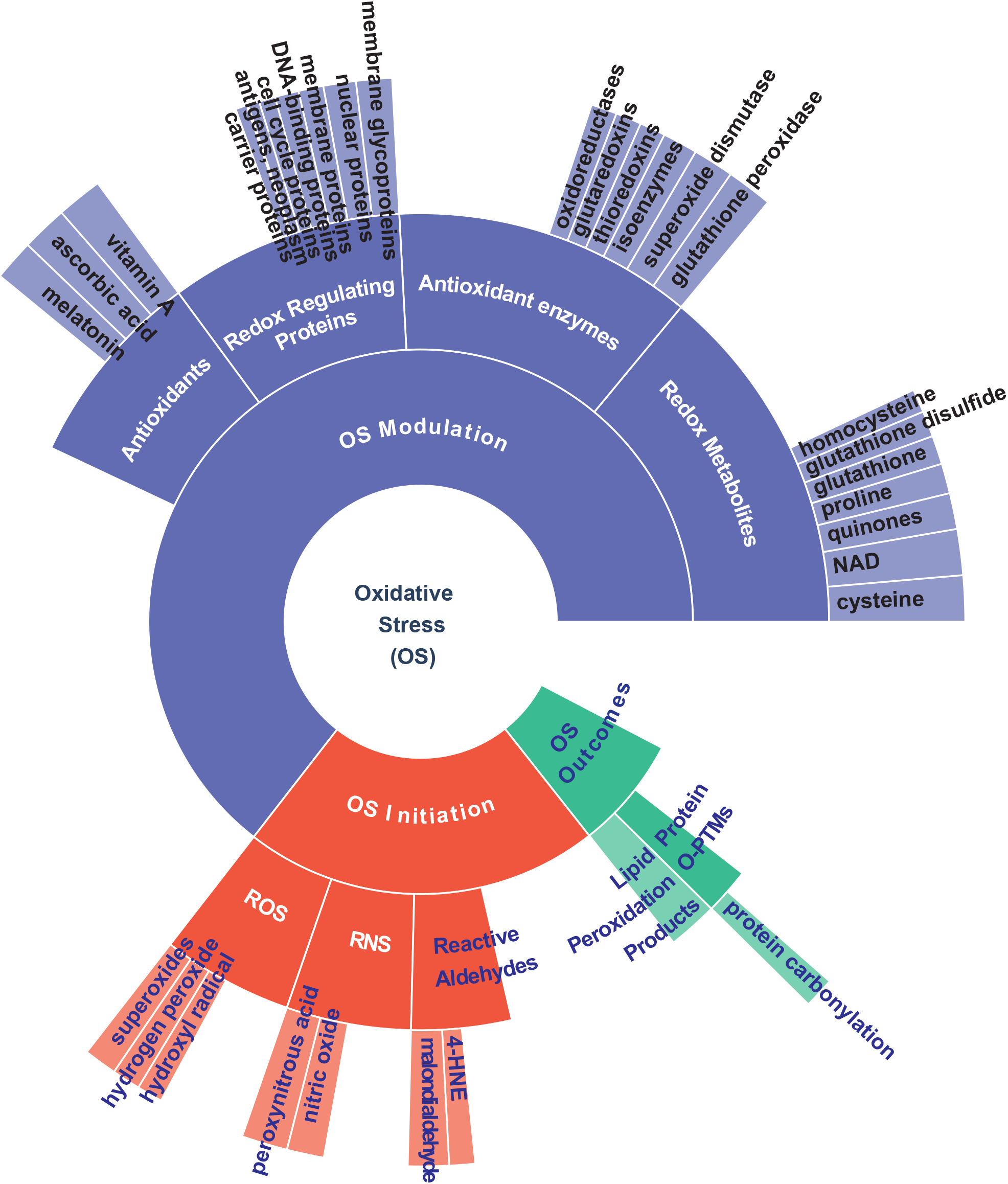
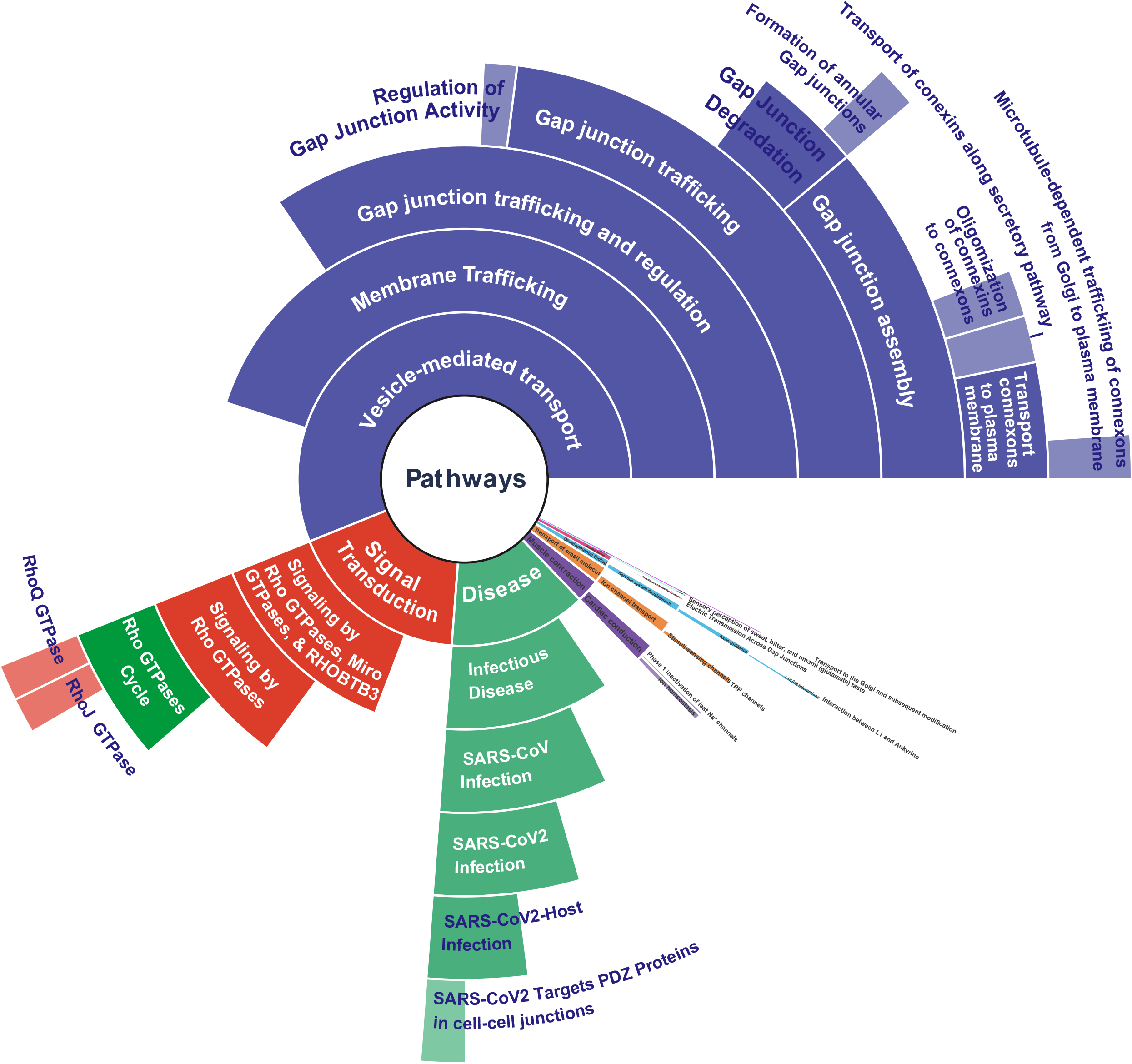
Significant OS molecules and pathways in an ARR-CCS cluster: This figure presents sunburst visualizations illustrating significant OS molecules (A) and pathways (B) linked to the proteins identified in the ARR and CCS clusters. Quantitative assessments of these OS molecules and pathways were conducted by calculating their respective scores based on the CaseOLAP scores of proteins within each CVD category. For OS molecules, scores were determined using the formula inside the parenthesis (OS molecule Score = Mean (Protein’s CaseOLAP score in each CVD category)). For pathways, the score was computed utilizing the formula inside the parenthesis (Pathway Score = Mean (Protein’s score in each CVD category *(1-p-value)). In both panels A and B, the arc thickness symbolizes the cumulative score for each molecule and pathway across the eight CVD categories. The outermost ring of the sunburst represents individual OS molecules (Panel A) and pathways (Panel B), with the arc width reflecting the score magnitude. The successive inner rings are organized hierarchically, with Panel A categorizing candidate molecules and Panel B sorting pathways based on their classification. An interactive version of these visualizations is available at https://caseolap.github.io/IonChannel/plots/os.html and https://caseolap.github.io/IonChannel/plots/os-cvd-pathways.html.

Our analysis shows that from a total of 75 OS molecules analyzed, 31 exhibit a significant association with the proteins under study. Dissected by OS phase, 5 molecules belong to the initiation phase, 23 to the modulation phase, and 3 to the outcome phase. All those OS molecules are provided in the supplemental **Table S4**.

Highlighted OS molecules in our study, such as hydroxyl radicals, superoxides, and hydrogen peroxide have been found in increased amounts in failing myocardium [38–40]. Experimental evidence has shown that increased concentration of these ROS molecules causes calcium overload in the cardiomyocytes by modulating the properties of Ca^2+^-regulating channels, ultimately causing contractile dysfunction and arrhythmias [40, 41]. For example, exposure to hydroxyl radicals causes Ca^2+^ overload in the cardiomyocytes by increasing the open probability of cardiac ryanodine receptors, which control Ca^2+^-release from the sarcoplasmic reticulum to the cytoplasm [42, 43]. Increased calcium influx through voltage-gated calcium channels is observed experimentally by brief exposure to hydrogen peroxide in ventricular myocytes [44, 45].

Conversely, antioxidants highlighted by our results (e.g., glutathione, glutathione peroxidase, superoxide dismutase, thioredoxins, vitamin a, and ascorbic acid) [46, 47], are essential for cardiovascular health [46]. Cells synthesize antioxidant compounds and enzymes to maintain redox homeostasis and mitigate ROS-induced damage. For example, superoxide dismutase utilizes superoxide to generate hydrogen peroxide, which catalase further metabolizes to water and oxygen [48]

##### 3.3.1.2. Significant Pathways in an ARR and CCS cluster

To elucidate the molecular mechanisms of protein clusters within the ARR and CCS categories, we collected associated molecular pathways using queries in KG. We scored relevant molecular pathways by using equation 2, which calculates pathway score by integrating both the protein associations and the significance levels (p-values) of the pathways.

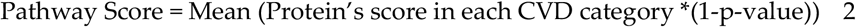

Initially, we collected all associated pathways tied to each protein. Since multiple proteins are associated with a single pathway, we utilized reverse mapping and collected all the proteins involved in the specific pathways. Next, we incorporated p-value of the pathway into the CaseOLAP score of associated proteins. We further took their means to create the final score for a pathway.

All the significant pathways in an ARR and CCS cluster are visualized in the interactive sunburst visualization (**Figure 6(B)**) following the hierarchical relationship of the pathways through the series of circles moving outwards according to their hierarchy. The pathway hierarchy is based on the Reactome knowledgebase. The significant pathways are distributed across the circles based on their hierarchy, where the thickness of the arc represents their collective scores in ARR and CCS, added to the total score of their descendants. There is a total of 50 significant pathways, and they are also provided in table format in the supplemental **Table S6**.

Some notable pathways highlighted by our exploration are Gap junction trafficking and regulation and their descendant pathways. The descendant pathways include Gap junction assembly, Gap junction degradation, Oligomerization of connexins into connexons, Transport of connexins along the secretory pathway, Transport of connexons to the plasma membrane, Microtubule-dependent trafficking of connexons from Golgi to the plasma membrane, and so on.

Gap junctions are clusters of intercellular channels connecting neighboring cells, facilitating the direct exchange of ions and small molecules. They are composed of connexins (6 transmembrane protein units) that are transported to the plasma membrane after oligomerizing into hexameric assemblies called connexons. The activity of these intercellular channels is regulated, particularly by intramolecular modifications such as phosphorylation which appears to regulate connexin turnover, gap junction assembly, and the opening and closure (gating) of gap junction channels. Excessive OS leads to reduced gap junction protein connexin (Cx43), a protein critical for normal cardiac conduction function. Reduced connexin levels may slow conduction and facilitate the proarrhythmic mechanism [49–51].

#### 3.3.2. KG Analysis: Link Prediction algorithm

We applied a link prediction algorithm to identify new biomedical knowledge from the patterns in the data [52]. In our case, we used link prediction to propose undiscovered relationships between (i) Ca^2+^-regulating proteins and OS molecules and (ii) Ca^2+^-regulating proteins and CVDs (**Figure 7**). It was implemented on our KG graph data from Neo4j’s Graph Data Science Library. The algorithm uses a graph’s topology to compute scores based on the closeness of a pair of nodes. These scores are then used to predict missing relationships. A higher score suggests a higher likelihood of an undiscovered node-node relationship.

**Figure 7.**
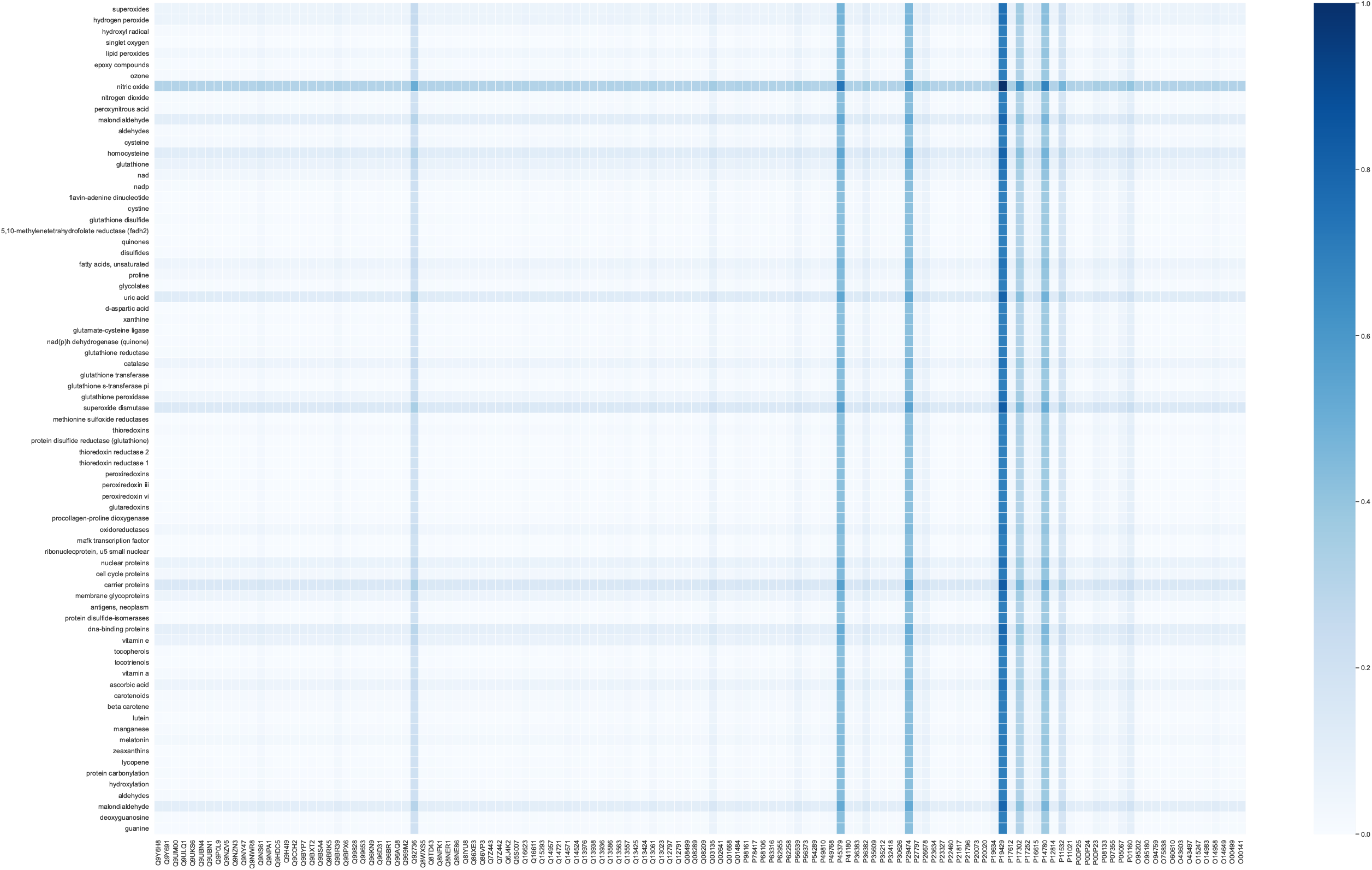
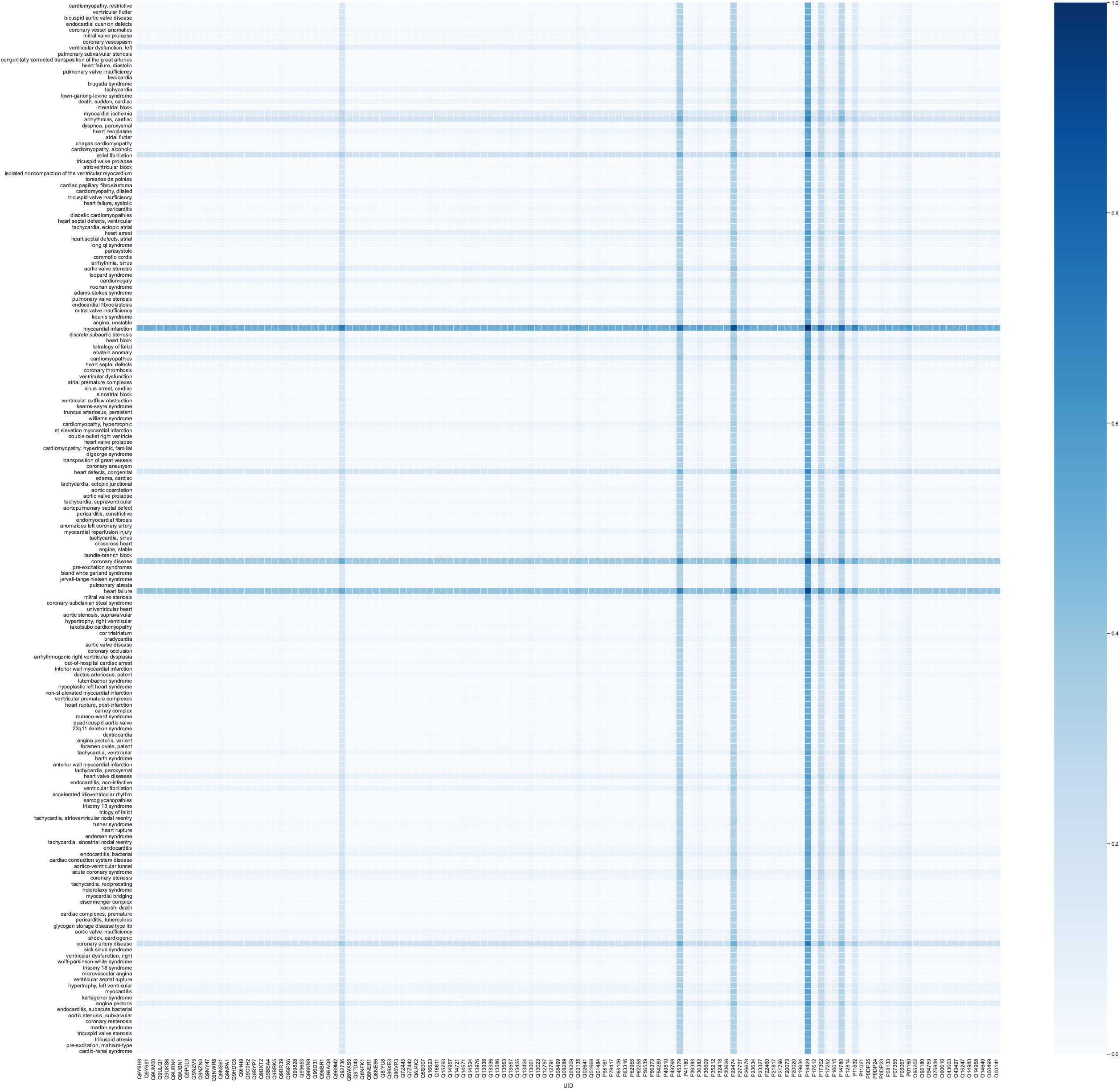
Link prediction analysis implementing graph algorithm: The heatmap depicts the predictive association scores derived from link prediction algorithms between OS molecules and Ca^2+^-regulating proteins (Panel A), as well as between Ca^2+^-regulating proteins and cardiovascular diseases (CVDs) (Panel B). The color intensity within the heatmap corresponds to the strength of the predicted association, with darker shades indicating a higher likelihood of connection between entities. Notably, proteins such as cardiac Troponin I, nitric oxide synthase, matrix metalloproteinase-9, and Gap junction alpha-1 protein, which play a role in calcium regulation, are identified with a high degree of probability as being associated with both OS molecules and CVD, underscoring their potential relevance in the underlying molecular mechanisms.

##### 3.3.2.1. Link Prediction between Ca^2+^-regulating proteins and OS MeSH descriptors

The link prediction algorithm ranked possible protein-OS relationships highlighting significant proteins-OS pairs. The top predicted protein-OS pair was cardiac troponin I, and nitric Oxide. In fact, it predicts that troponin I links with all OS molecules and that nitric oxide links with all proteins with a significant link prediction score (see **Figure 7(A)**). Both are essential and responsible for the heart’s excitation-contraction mechanism. Cardiac troponins are Ca^2+^-regulatory proteins accountable for the heart’s excitation-contraction mechanism [53–55]. NO influences the Ca^2+^-channels, facilitates the cGMP and PKG-dependent phosphorylation of troponin I, and attenuates myofilament response to calcium. Moreover, nitric oxide produced by endothelial cells regulates the excitation-contraction cycle of the heart by promoting vascular relaxation [56–59].

The other significant proteins were Matrix metalloproteinase-9, Nitric oxide synthase, endothelial, and Gap junction alpha-1 protein, and their association with Nitric Oxide is significantly higher than other OS MeSH descriptors. The table of the link prediction score is also provided in the supplemental **Table S7**.

##### 3.3.2.2. Link Prediction between Ca^2+^-regulating proteins and CVD MeSH descriptors

The link prediction analysis between proteins and CVD MeSH descriptors predicted the notable protein-CVD pair. The top predicted protein-CVD pair was myocardial infarction and Troponin I indicating a strong possible connection between them (see **Figure 7(B)**). The other top 4 proteins associated with myocardial infarction with higher link prediction scores were Nitric oxide synthase, endothelial, Matrix metalloproteinase-9, Gap junction alpha-1 protein and Ryanodine receptor 2. These proteins were also strongly associated with the OS MeSH descriptors (see **Figure 7(A)**) such as nitric oxide, carrier proteins, superoxide dismutase, and other CVD MeSH descriptors specifying the interconnection between the proteins, OS and CVDs. For example, our result highlighted a triangular association between myocardial infarction, Troponin I, and nitric oxide. The table of the link prediction score is also provided in the supplemental **Table S8**.

## 4. Discussion

Cardiac Ca^2+^-regulating proteins are critical for maintaining normal heart function via the cardiac excitation-contraction cycle. Emerging evidence has revealed that the overproduction of ROS such as hydroxyl radicals, superoxide, and hydrogen peroxide can interfere with the functionality, expression, and molecular pathways associated with Ca^2+^-regulating proteins. Such ROS-induced alterations in these proteins disturb both cytoplasmic and mitochondrial calcium homeostasis, leading to a deleterious feedback loop where imbalanced cellular calcium levels further amplify ROS production. This cycle is a contributing factor to the development of a variety of cardiac pathologies.

In this study, we utilize a text-mining approach combined with a knowledge graph to delineate the association of cardiac Ca^2+^-regulating proteins and oxidative stress in the context of 8 categories of cardiovascular disease. By integrating fragmented pieces of data pertaining to proteins, oxidative stress MeSH, and CVDs MeSH from an extensive corpus of PubMed articles, we sought to discover potential underlying connections that may have been overlooked. Our analysis provides insight into potential links between Ca^2+^-regulating proteins and oxidative stress across various CVDs, suggesting shared molecular pathways that may underlie these associations.

Our text mining analyses assume that the more that concepts are studied, the more biomedically relevant they are. In other words, the more often a Ca^2+^-regulating protein is mentioned in an accessible PubMed study with metadata annotating the OS and CVD categories in the study, the more biomedically relevant the protein, OS concept, and CVD are to one another. While this certainly does not hold perfectly, we believe that our strategy can meaningfully synthesize a vast amount of existing knowledge from disparate sources and draw new connections among the data.

## 5. Conclusions

Mutual interaction between the Ca^2+^-regulating proteins and OS has been associated with cellular processes in cardiovascular health and disease. In the present study, we investigated the impact of OS on cardiac Ca^2+^-regulating proteins with respect to the 8 major CVD categories by utilizing a text-mining algorithm combined with a knowledge graph. The key findings are 1) Ca^2+^-regulating proteins are distinctly associated with cardiac arrhythmias (ARR) and cardiac conduction system (CCS) diseases compared to other CVD categories. 2) 59 out of the 128 Ca^2+^-regulating proteins are found in OS-associated CVDs. 3) Cardiomyopathy is the highest-occurring disease category associated with mutual effect of OS and Ca^2+^-regulating proteins. 4) Utilizing a link prediction algorithm, hidden and possible relationships were emphasized among Ca^2+^-regulating proteins, OS molecules and CVDs. Our informatics study on the participation of Ca^2+^-regulating proteins in OS-associated cardiac diseases should help find novel early-detection tools and therapeutic strategies.

## Supplementary Materials

The following supporting information can be downloaded at: www.mdpi.com/xxx/s1, Table S1: CVDs MeSH Descriptors; Table S2: Oxidative Stress MeSH Descriptors; Table S3: Ca^2+^-regulating proteins; Table S4: Ca^2+^-OS-CVD interface; Table S5: OS molecules associated with proteins in ARR and CCS; Table S6: pathways associated with proteins in ARR and CCS; Table S7: OS-Protein Link prediction score; S8: CVD-protein Link Prediction Score

## Author Contributions

Conceptualization, P.P.; methodology, N.P., D.S., D.W., I.A., J.R., A.V., A.E.; software, N.P., D.S., I.A., J.R.; validation, N.P., D.S., D.W., and P.P; formal analysis, N.P., D.S., and D.W.; investigation, N.P., D.S., D.W., and P.P; resources, P.P.; data curation, N.P., D.S., and D.W.; writing—original draft preparation, N.P.; writing—review and editing, N.P., D.S., I.A., J.R., A.V., A.E., W.W., D.W., and P.P; visualization, N.P., D.S., and D.W.; supervision, P.P., and D.W.; project administration, N.P.; funding acquisition, P.P.

## Institutional Review Board Statement

Not applicable.

## Informed Consent Statement

Not applicable.

## Data Availability Statement

Not applicable.

## Acknowledgments

We thank Dr. Rajasekaran Namakkal-Soorappan and Dylan Steinecke for their support and helpful discussions.

